# The De Bruijn Graph Sequence Mapping Problem with Changes in the Graph

**DOI:** 10.1101/2024.02.15.580401

**Authors:** Lucas B. Rocha, Said Sadique Adi, Eloi Araujo

## Abstract

In computational biology, mapping a sequence *s* onto a sequence graph *G* poses a significant challenge. One possible approach to tackling this problem is to find a walk *p* in *G* that spells a sequence most similar to *s*. This challenge is formally known as the Graph Sequence Mapping Problem (`GSMP`). In this paper, we delve into an alternative problem formulation known as the De Bruijn Graph Sequence Mapping Problem (`BSMP`). Both problems have three variants: changes only in the sequence, changes in the graph, and changes in both the sequence and the graph. We concentrate on addressing the variant involving changes in the graph. In the literature, when this problem does not allow the De Bruijn graph to induce new arcs after changes, it becomes NP-complete, as proven by Gibney *et. al* [4]. However, we reformulate the problem by considering the characteristics of the arcs induced in the De Bruijn graph. This reformulation alters the problem definition, thereby enabling the application of a polynomial-time algorithm for its resolution. Approaching the problem with this arc-inducing characteristic is new, and the algorithm proposed in this work is new in the literature.

## I. INTRODUCTION

Mapping a sequence onto another for comparative purposes is a important task in computational biology. Typically, a comparison is made between one sequence and a reference sequence, acknowledged as a high-quality representation of a sequence set [9], [10]. However, the reference sequence is inherently biased as it only reflects a limited set of sequences and cannot encompass all potential variations. To address this bias, one approach is to represent multiple sequences through a more robust structure, such as a sequence graph or De Bruijn graph [6], [7], [13], [14].

The *sequence graph* is a graph such that each node is labeled with one or more characters and the *simple sequence graph* is one where each node is labeled with exactly one character [7]. In the *De Bruijn graph* [13], [14] of order *k*, each node is labeled with a distinct sequence of length *k* and there is an arc from one node to another if and only if there is an overlap of length *k* −1 from the suffix of the first to the prefix of second.

Informally, a walk *p* in a graph *G* is a sequence of connected nodes by arcs. Given a sequence graph *G*, a walk *p* in *G* can spell a sequence *s*^′^ by concatenating the characters associated with each node of *p*. The *Graph Sequence Mapping Problem* – `GSMP` consists of finding a walk *p* in a sequence graph *G* that spells a sequence as similar as possible to a given sequence *s*.

One of the first articles that addresses `GSMP` in more details was written by Amir *et. al*. in the article entitled *Pattern Matching in Hypertext* [1]. Navarro improved the results of this article and detailed these improvements in the article entitled *Improved Approximate Pattern Matching on Hypertext* [8]. For the approximate mapping, Amir *et. al*. were the first authors in the literature who identified an asymmetry in the location of the changes, showing the importance of understanding whether changes happen only in the pattern, only in the hypertext or in both. Considering the asymmetry identified by Amir *et. al*., the `GSMP` allows three variants:

1. allows changes only in the pattern when mapping the pattern in hypertext;
2. allows changes only in the hypertext when mapping the pattern in hypertext;
3. allows changes in the pattern and hypertext when mapping the pattern in hypertext.

Focusing exclusively on variant 1, it has been thoroughly studied in recent years by various authors, and this variant admits a polynomial-time algorithm. For example, Amir *et. al* [1] proposed a polynomial algorithm that runs in time *O*(*m*(|*V*| · log |*V*| + |*A*|)), and Navarro [8] improved it to *O*(*m*(|*V*| + |*A*|)). In the case of variants 2 and 3, Amir *et. al* proved that the respective problems are NP-complete when considering Hamming and edit distance, especially when the alphabet Σ has |Σ| ≥ |*V*| . In the work titled *On the Complexity of Sequence-to-Graph Alignment* [3], Jain *et. al* demonstrated that variants 2 and 3 are NP-complete even when the alphabet Σ has |Σ| ≥ 2.

The first time the `GSMP` was addressed using a De Bruijn graph as input was in the article entitled *Read Mapping on De Bruijn Graphs* [2]. In this work, Limasset *et. al*. propose the following problem, called here *De Bruijn Graph Sequence Mapping Problem* – `BSMP`: given a De Bruijn graph *G*_*k*_ and a sequence *s*, the goal is to find a path *p* in *G*_*k*_ such that the sequence *s*^′^ spelled by *p* have at most *d* differences between *s* and *s*^′^ with *d* ∈ℕ. The `BSMP` was proved to be NP-complete considering the Hamming distance, leading to the development of a seed-and-extended heuristic by the mentioned authors. Note that for the `BSMP` it does not make sense to talk about the three variants mentioned above since there are no node repetitions.

Considering that we extensively explored this issue in our recent paper titled *Heuristics for the De Bruijn Mapping Problem* [15], specifically focusing on variant 1 for the De Bruijn graph, our current focus is on investigating variant 2, which involves changes only in the graph structure. The `BSMP` was previously addressed in the context of walks in an article titled *On the Hardness of Sequence Alignment on De Bruijn Graphs* [4], where Gibney *et. al*. established its NP-completeness when changes occur in the graph.

Our current work is inspired by the study by Gibney *et. al* that demonstrated the NP-completeness of the problem for variant 2. However, we introduce a formulation of `BSMP` based on a specific characteristic of the De Bruijn graph, which involves inducing new arcs when a change is made to the graph. This formulation enables the application of a polynomial algorithm to solve the problem in this particular variant. In the study conducted by Gibney *et. al*, they do not account for this characteristic, resulting in the problem being NP-complete. In contrast, our study is pioneering in considering the induction of new arcs in the De Bruijn graph, making this polynomial algorithm groundbreaking for this problem.

## II. PRELIMINARIES

In this section, we describe some necessary concepts such as computational definitions and problem definition that are used in this paper.

### A. Sequence, distance, graphs and matching

Let Σ be an **alphabet** with a finite number of characters. We denote a sequence (or string) *s* over Σ by *s*[1]*s*[2] … *s*[*n*] in which each character *s*[*i*] ∈ Σ. We say that the **length** of *s*, denoted by |*s*|, is *n* and that *s* is a *n***-length** sequence.

We say that the sequence *s*[*i*]*s*[*i*+1] … *s*[*j*] is a **substring** of *s* and we denote it by *s*[*i, j*]. A substring of *s* with length *k* is a *k*-length sequence and also called *k***-mer** of *s*. For 1 ≤ j ≤ n in which *n* =| *s*|, a substring *s*[1, *j*] is called a **prefix** of *s* and a substring *s*[*j, n*] is called a **suffix** of *s*.

Given five sequences *s, t, x, w, z*, we define *st* the **concatenation** of *s* and *t* and this concatenation contains all the characters of *s* followed by the characters of *t*. If *s* and *t* are *n*- and *m*-length sequences respectively, *st* is a (*n*+*m*)-length sequence. If *s* = *xw* (*x* is a prefix and *w* is a suffix of *s*) and *t* = *wv* (*w* is a prefix and *z* is a suffix of *t*), we say the substring *w* is an **overlap** of *s* and *t*.

The **Hamming distance** *d*_*h*_ of two *n*-length sequences *s* and *t* is defined as

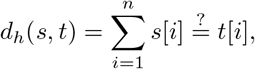

where 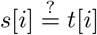 is equal to 1 if *s*[*i*] ≠ *t*[*i*], and 0 otherwise. In this context, we also say that *s* and *t* have *d*_*h*_(*s, t*) differences.

A **graph** is an ordered pair (*V, A*) of two sets in which *V* is a set of elements called **nodes** (of the graph) and *A* is a set of ordered pairs of nodes, called **arcs** (of the graph). Given a graph *G*, a **walk** in *G* is a sequence of nodes *p* = *v*_1_, …, *v*_*k*_, such that for each pair *v*_*i*_,*v*_*i*+1_ of nodes in *p* there is a arc (*v*_*i*_, *v*_*i*+1_) ∈ *A*. A **path** in *G* is a walk with no repeated nodes. Given a walk *p* = *v*_1_, …, *v*_*k*_,| *p*| = *k* −1 is the **length** of *p*. For graphs with costs associated with their arcs, the **cost** of a walk *p* is the sum of the cost of all arcs of all consecutive pairs of nodes (*v*_*i*_, *v*_*i*+1_) in *p*. A **shortest path** from *v*_1_ to *v*_*k*_ is one whose cost is minimum (a path of **minimum cost**).

An undirected graph *G* = (*V, A*) is called a **bipartite graph** *H* = (*V*_1_ ∪*V*_2_, *A*) if there is a partition of its nodes *V* into two disjoint sets *V*_1_ and *V*_2_, such that for each edge (*u, v*) ∈ *A, u* ∈ *V*_1_ and *v* ∈ *V*_2_. A bipartite graph is called a **complete bipartite graph** if there is an edge from every vertex *u* ∈ *V*_1_ to every vertex *v* ∈ *V*_2_.

A **matching** *E* in *H* consists of a set of *m* edges of *H* that do not share common nodes. The **cost of the matching** *E* is *C*(*E*) =∑ _*e ∈ E*_ *z*(*e*), where *z*(*e*) is the cost of an edge *e* = {*x*, y} ∈ *A*. A matching is said to be of **minimum cost** if there is no matching with a lower cost. A matching *E* is **maximum** if there is no other matching larger than *E*, i.e., if there is no matching *E*^′^ such that |*E*^′^| > |*E*| . Given a set E with all maximum matchings, a matching *E* ∈ E is said to be **maximum and of minimum cost** if there is no matching *E*^′^ such that *C*(*E*^′^) < C(*E*).

Given a bipartite graph *H* = (*V*_1_ ∪ *V*_2_, *A*), find a minimum-cost matching *E*. This problem is extensively studied in the literature and can be solved using, for example, Edmonds’ algorithm (with a time complexity of *O*((|*V*_1_|+|*V*_2_|)^3^)), Hopcroft-Karp algorithm (with a time complexity of 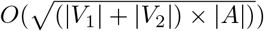, and the Hungarian algorithm (with a time complexity of *O*((|*V*_1_| + | *V*_2_|)^3^)) [11], [12], [17].

A **sequence graph** is a graph (*V, A*) with a sequence of characters, built on an alphabet Σ, associated with each of its nodes. A **simple sequence graph** is a graph in which each node is labeled with only one character. Given a set *S* = *r*_1_, …, r_*m*_ of sequences and an integer *k* ≥2, a **De Bruijn graph** is a graph *G*_*k*_ = (*V, A*) such that:

- *V* = {*d* ∈ Σ^*k*^| such that *d* is a substring of length *k* (*k*-mer) of *r* ∈ *S* and *d* labels only one node};
- *A* = {(*d, d*^′^)| the suffix of length *k* − 1 of *d* is a prefix of *d*^′^}.

Note that we can define the De Bruijn graph just by its set of nodes and the arcs are induced for each pair of *k*-mer *d* and *d*^′^ whose *k* −1 length prefix of d is equal to the *k* −1 length suffix of *d*^′^.

In this paper, informally for readability, we consider the node label as node. Given a walk *p* = *v*_1_, *v*_2_, …, *v*_*n*_ in a De Bruijn graph *G*_*k*_, the **sequence spelled** by *p* is the sequence *v*_1_*v*_2_[*k*] … *v*_*n*_[*k*], given by the concatenation of the *k*-mer *v*_1_ with the last character of each *k*-mer *v*_2_, …, *v*_*n*_. For a walk *p* = *v*_1_, *v*_2_, …, *v*_*n*_ in a simple sequence graph *G*, the sequence spelled by *p* is *v*_1_*v*_2_ … *v*_*n*_. A **mapping** of a sequence *s* onto a simple sequence graph or a De Bruijn graph *G* is a walk *p* in *G* whose editing cost between *s* and the sequence spelled by *p* is minimum.

Given the definitions above, we state the following problem for simple sequence graphs (`GSMP`) and for De Bruijn graphs (`BSMP`) when we have changes only in the sequence, respectively:

*Problem 1 (****Graph Sequence Mapping Problem –*** `GSMP`): Given a *m*-sequence *s* and a simple sequence graph *G*, find a mapping of *s* onto *G*.

*Problem 2 (****De Bruijn Graph Sequence Mapping Problem –*** `BSMP`): Given a *m*-sequence *s* and a De Bruijn graph of order *k, G*_*k*_, find a mapping of *s* onto *G*_*k*_.

## III. FORMULATING THE DE BRUIJN MAPPING PROBLEM

We address a variant of the SEQUENCE MAPPING PROBLEM IN DE BRUIJN GRAPHS, which considers changes exclusively in the graph. Unlike Gibney *et. al* [4], who defined a problem without allowing the graph structure to change and proved its NP-completeness, in this work, we allow the graph structure to be altered. In summary, the main difference between this study and the work conducted by the authors is whether or not we allow the alteration of arcs when we change the vertex labels. For the variant studied in this chapter, we formulate the problem by allowing the induction of new arcs in a De Bruijn graph whenever a vertex label is modified, and this flexibility allows us to find solutions in polynomial time.

Given that the De Bruijn graph allows the induction of arcs through its *k*-mers, we consider the definitions in the next two sections to redefine the problem taking into account this characteristic of induced arcs.

### A. The edition of a De Bruijn graph

Let *s* be a sequence and *G*_*k*_ a set of nodes labeled by *k*-mers representing a De Bruijn graph (the edges are implicit). We define *k*(*s*) as the set of *k*-mers in *s* and assume that *m* = |*k*(*s*) |≤ |*G*_*k*_| = *n*.

An **edit** *E* with *j* operations in *G*_*k*_ with respect to *s*, or simply edit *E* in *G*_*k*_, consists of replacing the labels of a sequence *v*_1_, *v*_2_, …, *v*_*j*_ of distinct nodes in *G*_*k*_ with a sequence *u*_1_, …, u_*j*_ of distinct *k*-mers in *k*(*s*) (each label *v*_*i*_ is replaced by *u*_*i*_), such that after the edit, there is a walk in the resulting De Bruijn graph that induces *s*. The graph obtained after the edit *E* is denoted by 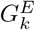. Note that every *k*-mer in *k*(*s*) is a label of some vertex in 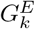. The **cost of the edit** *E* is

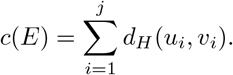

Since every *k*-mer in *k*(*s*) is a label of some vertex in 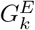, if *j* < *m*, it means that *m* − *j k*-mers *u*_*j*+1_, u_*j*+2_, …, u_*m*_ in *k*(*s*) are labels of *m − j* nodes *v*_*j*+1_, *v*_*j*+2_, …, *v*_*m*_ in *G*_*k*_ with *d*_*H*_ (*u*_*i*_, *v*_*i*_) = 0 for each *i* = *j* + 1, *j* + 2, …, *m*, implying that replacing the labels of *v*_1_, *v*_2_, …, *v*_*m*_ with *u*_1_, u_2_, …, u_*m*_ is an edit *E*^′^ of the same cost as the edit *E*, that is,

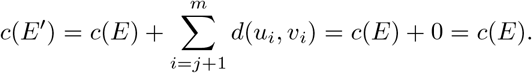

We then assume, without loss of generality, that every edit has exactly *m* operations. The set ℰ is defined as the set of all edits in *G*_*k*_. An **optimal edit** is then an edit of minimal cost, and we denote the cost of an optimal edit in *G*_*k*_ with respect to *s* by *D*_*H*_ (*s, G*_*k*_) = min_*E*∈*E*_ c(*E*). To exemplify these definitions, consider, for example, the sequence *s* = AACCAACCA and the De Bruijn graph *G*_3_ = `AAC`, `ACC`, `CAG`, `TTT` . In this case, we have *k*(*s*) = `AAC`, `ACC`, `CCA` and `CAA`, and Figure 1 shows examples of an edit and optmial edit in *G*_3_ for this case.

**Figure 1.**
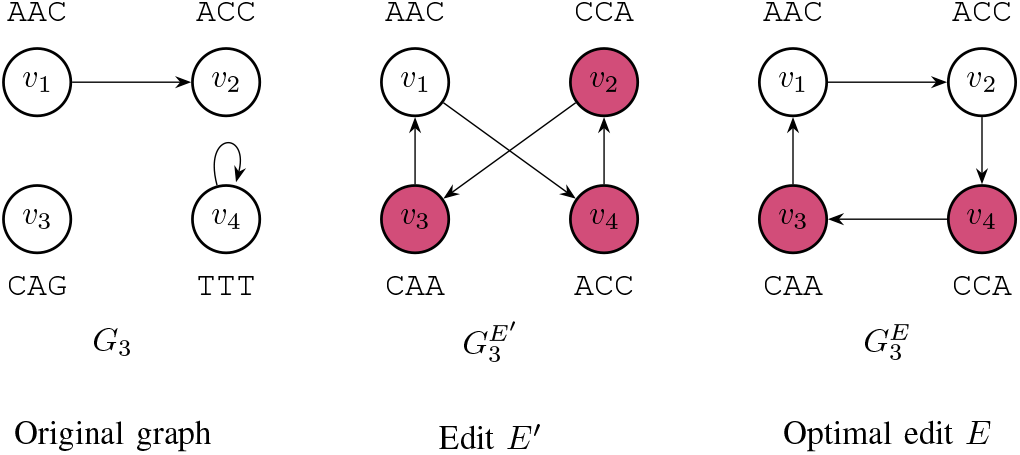
Example of an edit *E^′^* in *G*_3_ for *s* =AACCAACCA that replaces the labels of *v*_2_, *v*_3_, and *v*_4_. The cost of *E*^′^ is *c*(*E*^′^) = 5, reflecting the cost to replace the label `ACC` with `CCA`, `CAG` with `CAA`, and the label `TTT` with `ACC`. Besides the edit *E*^′^, we have an example of an optimal edit *E* in *G*_3_ for the same *s* = AACCAACCA that replaces the labels of *v*_3_ and *v*_4_. The cost of *E* is *c*(*E*) = 4, reflecting the cost to replace the label `CAG` with `CAA` and the label `TTT` with `CCA`. As a result of the edit *E*, we have the De Bruijn graph 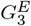 . Note that the walk *p* = *v*_1_, *v*_2_, *v*_4_, *v*_3_, *v*_1_, *v*_2_, *v*_4_ in 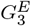 induces the sequence *s*.

### B. Bipartition and Matching

Next, we describe the creation of a bipartite graph from *k*(*s*) and a De Bruijn graph *G*_*k*_ (represented only by its nodes) and the relationship between a minimum cost maximum matching and the optimal edit in *G*_*k*_ with respect to *s*. For two sets *A* and *B*, we denote by *A* ×*B* the set formed by all nodes of *A* and *B* such that *u* ∈ *A* and *v* ∈ *B*, that is, *A* ×*B* = *u, v* : *u* ∈ *A* and *v* ∈ *B*. Consider *H* = (*k*(*s*) ∪*G*_*k*_, *k*(*s*) ×*G*_*k*_) a complete bipartite graph where each vertex in *G*_*k*_ is adjacent to each of the *k*-mers in *k*(*s*). The cost of an edge *u, v* in *H* is given by *d*_*H*_ (*u, v*).

A maximum matching, or simply matching *M*, in *H* has *m* edges and has a cost

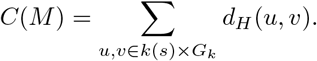

Note that since |*k*(*s*)|≤|*G*_*k*_|, this matching covers all nodes of *k*(*s*). We define here that an **optimal matching** *M*^*^ is a maximum matching of minimum cost.

Let *M* = *u*_1_, *v*_1_, …, u_*m*_, *v*_*m*_ be a matching in *H* = (*k*(*s*) ∪ G_*k*_, k(s)×*G*_*k*_). The edit *E*_*M*_ with *m* operations corresponding to the matching *M* is the edit where each edge *u, v* of the matching (*u* ∈ k(s) and *v*∈ *G*_*k*_) corresponds to an operation of replacing the label of *v* with *u*. We say that *M* transforms *G*_*k*_ into 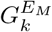. Note that by the definition of edit and matching,

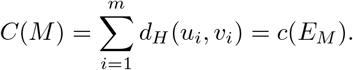

Similarly, the operation of replacing each label of *v* with *u* in an edit *E* corresponds to an edge in a matching, denoted by *M*_*E*_, in *H* = (*k*(*s*) ∪ *G*_*k*_, *k*(*s*) × *G*_*k*_), and note again that by the definition of edit and matching,

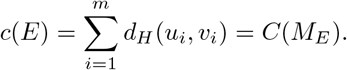

An example of this complete bipartite graph, matching, and the cost of an edit is shown in Figure 2, for the De Bruijn graph *G*_3_ = `ACT`, C`TG`, A`CG`, T`TT` and *k*(*s*) = `ACT`, C`TG`, T`GC`, G`CG` obtained from the sequence *s* =AACCAACCA. In this example, the cost of each edge *e* = *u, v*, where *u* ∈ *k*(*s*) and *v* ∈ *G*_*k*_, is determined by *c*(*e*) = *d*_*H*_ (*u, v*).

**Figure 2.**
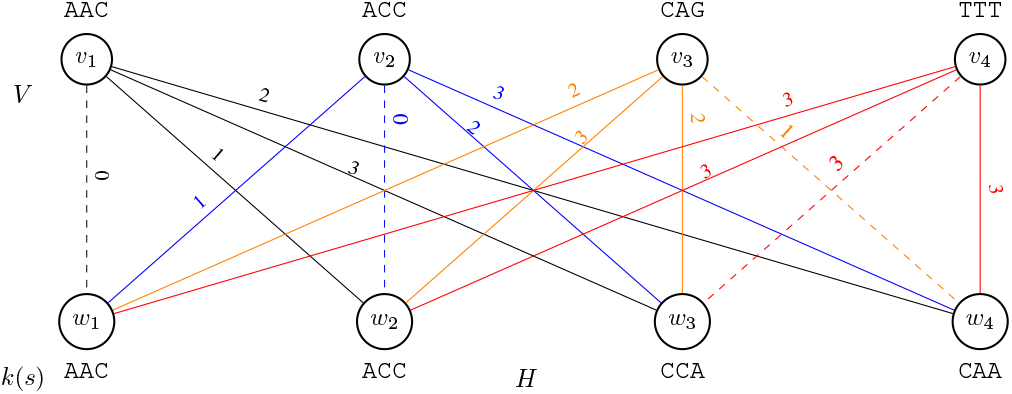
Example of a complete bipartite graph *H* = (*k*(*s*)∪ *G*_*k*_, *k*(*s*) ×*G*_*k*_) for the De Bruijn graph *G*_3_ = { A`AC`, `ACC`, `CAG`, `TTT`} and *k*(*s*) = {`AAC`, `ACC`, `CCA`, `CAA`}, obtained from the sequence *s* =AACCAACCA. Each edge *e* = {*u, v*} ∈*A* has an associated cost *c*(*e*) = *d*_*H*_ (*u, v*). The dotted edges represent an optimal matching *M* ^*^ = {{*u*_1_, *v*_1_},{ *u*_2_, *v*_2_}, {*u*_3_, *v*_4_}, {*u*_4_, *v*_3_}, with a total cost of 4. The cost of the edit *E*_*M*_* is equal to 4, considering the alteration of the *k*-mer `CAG` at vertex *v*_3_ to `CAA` and the *k*-mer `TTT` at vertex *v*_4_ to `CCA`. The colors on the edges serve to facilitate the visualization of the graph.

### C. Theorem

The following theorem is used to describe an algorithm that finds an optimal edit in *G*_*k*_ with respect to *s*.

*Theorem 1:* Let *s* be a sequence, and *G*_*k*_ be a De Bruijn graph. If *M*^*^ is an optimal matching in *H* = (*k*(*s*)∪ *G*_*k*_, *k*(*s*)× *G*_*k*_), then *C*(*M*^*^) = *D*_*H*_ (*s, G*_*H*_), and *E*_*M*_* corresponds to an optimal edit in *G*_*k*_ with respect to *s*.

**Proof**. The edit *E*_*M*_* (corresponding to the matching *M*^*^ in *H*) transforms *G*_*k*_ into 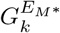and has a cost *c*(*E*_*M*_*) = *C*(*M*^*^), thus *D*_*H*_ (*s, G*_*k*_) ≤*c*(*E*_*M*_*) = *C*(*M*^*^). On the other hand, a matching *M*_*E*_* (corresponding to the optimal edit *E*^*^ in *G*_*k*_) in *H* has a cost *C*(*M*_*E*_*) = *D*_*H*_ (*s, G*_*k*_), which implies that *C*(*M*^*^)≤ *C*(*M*_*E*_*) = *D*_*H*_ (*s, G*_*k*_).

Therefore, *C*(*M*^*^) = *c*(*E*_*M*_*) = *D*_*H*_ (*s, G*_*H*_), implying that *C*(*M*^*^) = *D*_*H*_ (*s, G*_*k*_) and consequently, the edit *E*_*M*_* corresponds to an optimal edit in *G*_*k*_ with respect to *s*. _□_

### D. Algorithm

Theorem 1 ensures that an algorithm to find an optimal matching in *H* can be used to find an optimal edit in *G*_*k*_, which is precisely what the following algorithm does. Given a De Bruijn graph *G*_*k*_ (represented solely by its set of nodes) and a sequence *s*, the proposed algorithm is called DE BRUIJN GRAPH MAPPING TOOL WITH GRAPH CHANGES – `BMTC`, and it follows these steps. Initially, the set *k*(*s*) is determined. Then, the complete bipartite graph *H* = (*k*(*s*) ∪ G_*k*_, k(s) × *G*_*k*_) is created, assigning a cost *d*_*H*_ (*u, v*) to each edge *u, v* with *u* ∈ k(s) and *v* ∈ G_*k*_, as specified in Algorithm **??** (BIPARTIDE). The Hungarian algorithm [17] is applied to the graph *H* to find a maximum matching of minimum cost *M*^*^. These steps to solve the sequence mapping problem in De Bruijn graphs with changes are detailed in the pseudocode shown in Algorithm 1, and the algorithm returns *M*^*^ and the cost of the optimal edit.

#### Algorithm 1

DE BRUIJN GRAPH MAPPING TOOL WITH GRAPH CHANGES – `BMTC`

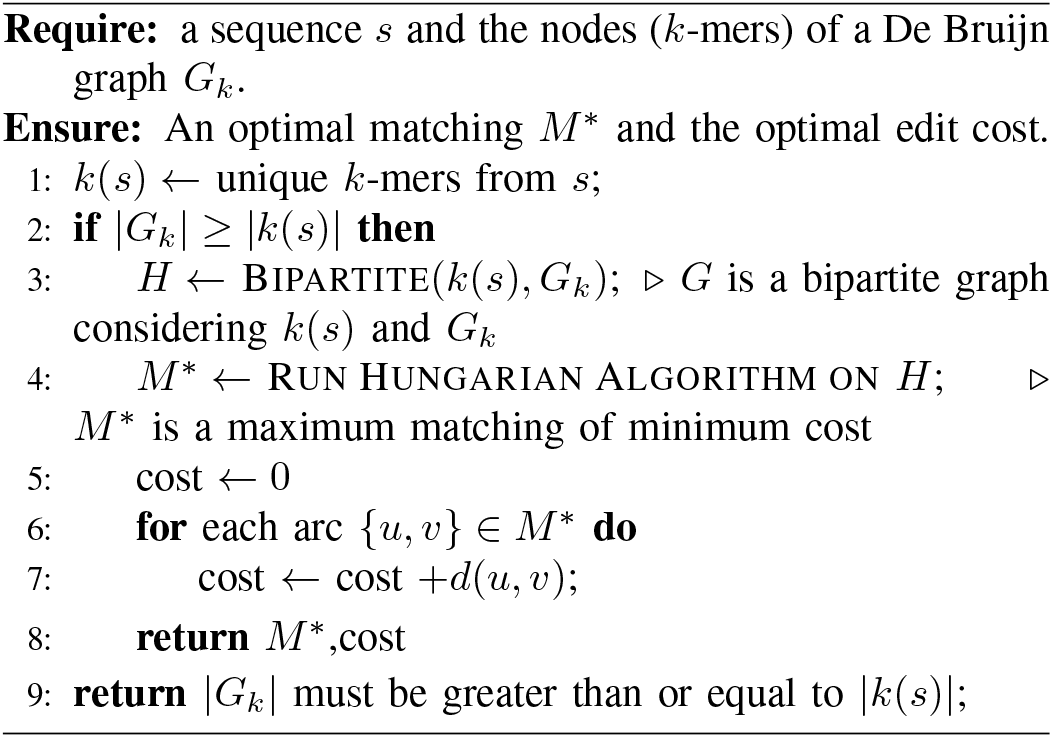

#### Algorithm 2

BIPARTITE

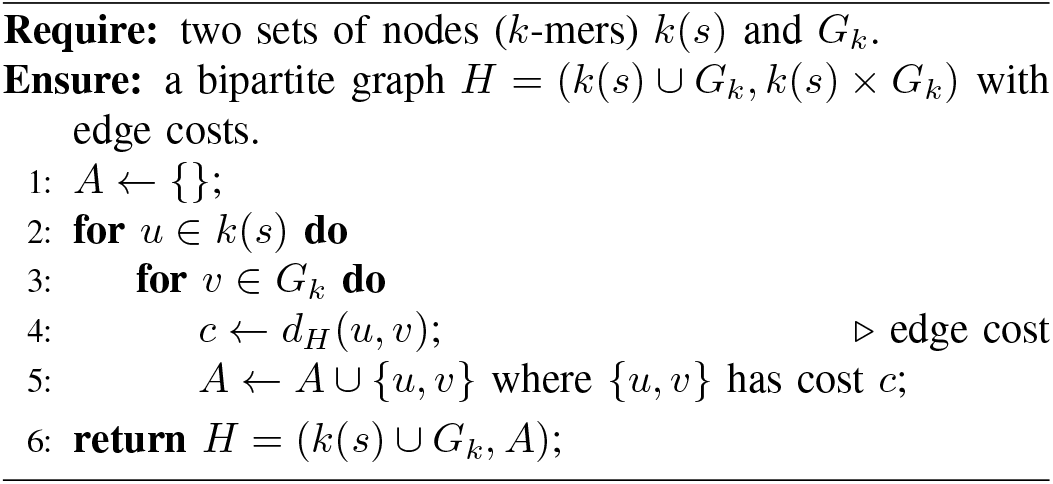

To determine the time complexity of Algorithm 1, consider *m* = |*k*(*s*)| ≤ |*G*_*k*_| = *n* and |*s*| = l. Thus, the time complexity for identifying all the *k*-mers in line1 is Θ(l *k*). In line3, creating the bipartite graph and computing the cost of the edges has a time complexity of *O*(*n m k*) because for each of the *n* nodes *v* of *G*_*k*_, we compute for each of the *m k*-mers *u* of *k*(*s*) the cost *d*_*H*_ (*u, v*), consuming time Θ(*k*). The execution of the Hungarian algorithm in line4 has a time complexity of *O*((*n* + *m*)^3^). Finally, iterating through all the edges of the matching and calculating the edit cost in lines6– 7 is *O*(*m*). Therefore, the total time complexity is given by Θ(l· *k*) +*O*(*n* ·*m*· *k*) +*O*((*n*+*m*)^3^) +*O*(*m*) = *O*((*n*+*m*)^3^ + (*n*· *m* ·*k*) + *m* + (l· *k*)) = *O*(*n*^3^ + l) since *k* is less than the number of nodes in the graph and *m* ≤ *n*. The implementation is available on github: github.com/LucasBarbosaRocha.

## IV. TESTS

In these tests, the objective is to use the developed tool named `BMTC` and evaluate the cost of modifications made in the graph, as well as the execution time of the tool. As discussed earlier, our proposal of `BMTC` is pioneering, as modifications in the graph induce new arcs, and therefore, there is no similar tool in the literature for comparison. In these tests, we used sequences with an average length of 5566 to map onto the graph, and De Bruijn graphs with *k*-mers ranging from 1464 to 11033. To assess `BMTC`, we performed the following operation in each test:

- Given all *k*-mers of the sequence *s* to be mapped onto the graph;
- Build a De Bruijn graph with these *k*-mers;
- To edit 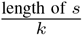 *k*-mers in the graph (note that this step may reduce the number of *k*-mers in the graph, as a modification can generate an already-existing *k*-mer);
- Run `BMTC`.

Given the previous steps, we have control over how many *k*-mers were altered in the graph, considering that a single modification was made to each *k*-mer in the graph, resulting in 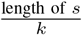 modifications in the graph. When executing `BMTC`, we obtain the pairing cost, and as expected, the cost is at most 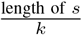. In these tests, we ran `BMTC` with *k* values equal to 10, 20, and 30. The results are presented in Table I. For each *k* value, we conducted 100 tests.

**Table I.**
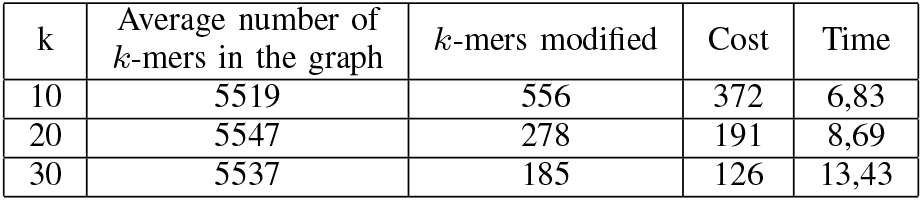
AVERAGE COSTS OBTAINED WITH THE BMTC TOOL AND THE AVERAGE EXECUTION TIME (IN SECONDS): TESTS CONDUCTED WITH MAPPED SEQUENCES OF AN AVERAGE LENGTH OF 5566.

| k | Average number of $k$-mers in the graph | $k$-mers modified | Cost | Time |
| --- | --- | --- | --- | --- |
| 10 | 5519 | 556 | 372 | 6,83 |
| 20 | 5547 | 278 | 191 | 8,69 |
| 30 | 5537 | 185 | 126 | 13,43 |

For *k* = 10, the number of *k*-mers in the graph varies between 1482 and 10933, with an average of 5519 *k*-mers. For *k* = 20, the number of *k*-mers in the graph varies between 1474 and 11043, with an average of 5547 *k*-mers. As for *k* = 30, the number of *k*-mers in the graph varies between 1464 and 11033, with an average of 5537 *k*-mers. It is observed in the table that, based on the average length of the sequence 5566, for *k* = 10, we have an average of 556 modified *k*-mers, with the value returned by `BMTC` being 372 (on average). For *k* = 20, we have 278 modified *k*-mers, with an average cost of 191 returned by `BMTC` for making modifications to the graph. As for *k* = 30, we have 185 modified *k*-mers, with an average cost of 126 for the modifications.

In the tests, we observed that the application executes in seconds, and it is also noted that increasing the value of *k* results in an increase in execution time. However, this application demonstrates promising performance, and the adopted strategy proves effective in executing modifications in the graph.

## V. DISCUSSION, PERSPECTIVES AND CONCLUSIONS

In this paper, we delve into the De Bruijn Problem Mapping with changes in the graph. We explore the work of Gibney *et. al* [4], which demonstrated the NP-completeness of the problem for De Bruijn graphs. In this work, we allow a change in the graph not only to modify the labels of the nodes but also the topology of the graph. The fundamental distinction lies in the flexibility allowed, exploring the induction of new arcs after a modification in the De Bruijn graph. While Gibney *et. al* focus on modifications that preserve the fundamental structure, this work considers changes that can induce new arcs, enabling polynomial-time solutions.

Considering this feature of inducing new arcs, we redefine the problem, introducing concepts such as the *s-transformation* of a De Bruijn graph and the *Bipartition and matching between two sets of k-mers*. We develop an algorithm called `BMTC`, which utilizes the Hungarian algorithm to find a maximum-cost minimum matching in a bipartite graph, resulting in a modified set of nodes for the De Bruijn graph.

The proof that the algorithm works is based on lemmas establishing the relationship between the graph transformation and matchings in the bipartite graph. The theorem demonstrates that the cost of the maximum matching found in the bipartite graph is equal to the Hamming distance between the given sequence and the original graph. `BMTC` offers an innovative approach to the problem, allowing changes in the De Bruijn graph that may result in new arcs, proving advantageous for finding polynomial-time solutions.

It is noteworthy that the literature still lacks depth in this aspect. The most recent work by Gibney *et. al* [[4]] addresses the problem but in a different manner. By introducing the possibility of inducing new arcs in case of changes in the De Bruijn graph, this work proposes a polynomial-time solution to the problem. However, further exploration of the potential practical applications of this solution is still needed.

## ACKNOWLEDGMENT

This research was supported by CAPES and UFMS.

